# SPINT1 regulates the aggressiveness of skin cutaneous melanoma and its crosstalk with tumor immune microenvironment

**DOI:** 10.1101/611145

**Authors:** Elena Gómez-Abenza, Sofía Ibáñez-Molero, Diana García Moreno, Inmaculada Fuentes, Leonard I. Zon, Maria C. Mione, María L. Cayuela, Chiara Gabellini, Victoriano Mulero

## Abstract

Skin cutaneous melanoma (SKCM) is the deadliest form of skin cancer and while incidence rates are declining for most cancers, they have been steadily rising for SKCM worldwide. Serine protease inhibitor, kunitz-type, 1 (SPINT1) is a type II transmembrane serine protease inhibitor that has been shown to be involved in the development of several types of cancer. We report here a high prevalence of *SPINT1* genetic alterations in SKCM patients and their association with altered tumor immune microenvironment and poor patient survival. We used the unique advantages of the zebrafish to model the impact of SPINT1 deficiency in early transformation, progression and metastatic invasion of SKCM. Our results reveal that Spint1a deficiency facilitates oncogenic transformation, regulates the tumor/immune microenvironment crosstalk, accelerates the onset of SKCM and promotes metastatic invasion. Notably, Spint1a deficiency is required at both cell autonomous and nonautonomous levels to enhance invasiveness of SKCM. These results suggest the relevance of clinical intervention on this signaling pathway for precision SKCM medicine.

**Summary statement:** A zebrafish model shows that Spint1a deficiency facilitates oncogenic transformation, regulates the tumor/immune microenvironment crosstalk, accelerates the onset of SKCM, and promotes metastatic invasion in cell autonomous and non-autonomous manners.

## Introduction

Skin cutaneous melanoma (SKCM) originates from melanocytes, neural-crest derived pigment-producing cells located in the epidermis, where their major function is to protect keratinocytes from UV-induced DNA damage (Wellbrock and Arozarena, 2016). The malignant transformation of melanocytes generates this fatal form of skin cancer with a complex multigenic etiology that becomes extremely difficult to treat once it has metastasized. SKCM is the deadliest form of skin cancer (75% of deaths related to skin cancer) and it is common in the Western world. Indeed, its global incidence is 15–25 per 100,000 individuals (Schadendorf and Hauschild, 2014). While incidence rates are declining for most cancers, they have been steadily rising for SKCM worldwide (van Rooijen et al., 2017). Early detection is fundamental, since localized, early stage SKCM can be surgically excised with little chance of recurrence with a 98.2% of patient survival rate after 5 year survival as reported by The Surveillance, Epidemiology, and End Results (SEER) (NIH, 2019). Metastatic SKCM, however, is still an often fatal disease with a 5-year survival rate of 15-20% (van Rooijen et al., 2017).

SKCM is one of the most recurrent types of cancer and its genetic heterogeneity has led in recent years to join forces to determine SKCM causes and develop effective therapies. Transformation of melanocytes into primary and then metastatic SKCM requires a complex interplay of exogenous and endogenous events (Schadendorf et al., 2015). More than 50% of the tumors originate from normal skin rather than from dysplastic nevi, suggesting that SKCM not only appears to be due to the transformation of mature melanocytes, otherwise it may arise from a malignant transformation of melanocytic progenitors (Hoerter et al., 2012) which sustain cancer development. In this way, the identification of SKCM initiating cells is really important to devising methods for early detection and eradication of SKCM (Kaufman et al., 2016; Santoriello et al., 2010). Moreover, SKCM stem cell populations have been characterized and associated with tumor progression, immunoevasion, drug resistance and metastasis (Nguyen et al., 2015).

Inflammation can play a key role in cancer, from initiation of the transformed phenotype to metastatic spread. Nevertheless, inflammation and cancer have a profound yet ambiguous relationship. Inflammation (especially chronic inflammation) has protumorigenic effects, but inflammatory cells also mediate an immune response against the tumor and immunosuppression is known to increase the risk for certain tumors (Shalapour and Karin, 2015). Nowadays, skin cancers are also attributed to chronically injured or non-healing wounds and scars or ulcers that occur at sites of previous burns, sinuses, trauma, osteomyelitis, prolonged heat and chronic friction. The incidence of malignancy in scar tissues is 0.1–2.5 % and it is estimated that underlying infections and inflammatory responses are linked to 15–20% of all deaths from cancer worldwide (Maru et al., 2014). Furthermore, chronic inflammation contributes to about 20% of all human cancers (Tang and Wang, 2016).

Serine protease inhibitor, kunitz-type, 1 (SPINT1), also known as hepatocyte growth factor activator inhibitor 1 (HAI1), is a type II transmembrane serine protease inhibitor that plays a crucial role in the regulation of the proteolytic activity of both suppression of tumorigenicity 14 (ST14), also known as Matriptase-1 (Benaud et al., 2001; Lin et al., 1999; Tseng et al., 2008), and Hepatocyte growth factor activator (HGFA) (Shimomura et al., 1997). The functional linkage between ST14 and SPINT1 has important implications for the development of cancer. ST14 activity, which is only partially opposed by endogenous SPINT1, causes increased sensitivity to carcinogens and produces spontaneous tumorigenesis in the skin of keratin-5-matriptase transgenic mice, while increased epidermal SPINT1 expression fully counteracts the oncogenic effect of ST14 (List et al., 2005). Furthermore, the expression of ST14 has been demonstrated to be up-regulated in various human cancer histotypes such as breast, cervix, ovaries, prostate, esophagus and liver cancers (List, 2009).

The close functional relationship between ST14 and SPINT1 was also observed in a zebrafish model of skin inflammation, carrying a hypomorphic mutation of *spint1a*. Indeed epidermal hyperproliferation and neutrophil infiltration observed in mutant zebrafish larvae are both rescued by *st14a* gene knock-down, suggesting a novel role for the SPINT1-ST14 axis in regulating inflammation (Carney et al., 2007; Mathias et al., 2007). Given the unique advantages of the zebrafish model for *in vivo* imaging and the strong correlation between alterations of Spint1a-St14a levels with tumor progression, the *spint1a* mutant zebrafish represents an attractive model to study the role of SPINT1 and chronic inflammation in SKCM.

Our results support the human data that show that genetic alterations of SPINT1 correlated with a poor prognosis of SKCM patients and provide evidence that SPINT1 expression positively correlated with tumor macrophage infiltration, but not neutrophils. In line with these clinical data, we show that Spint1a deficiency enhances at both cell autonomous and non-autonomous levels cell dissemination of SKCM in zebrafish models by promoting tumor dedifferentiation and altered immune surveillance.

## Results

### SPINT1 genetic alterations are associated with poor prognosis of SKCM patients and altered tumor immune microenvironement

To study the impact of *SPINT1* in promoting SKCM progression and aggressiveness, an *in silico* analysis of human SKCM samples of the TCGA cohort was performed. This analysis revealed that genetic alterations occurred in 10% of SKCM patients; a relevant percentage comparing with major SKCM driven oncogenes and tumor suppressors (**Figure 1A**). Among these genetic alterations, an increased mRNA level was the most prevalent alteration (7%), while 1.9% missense mutations of unknown significance and 1.9% deep deletions were also observed. Notably, these genetic alterations of *SPINT1* significantly correlated with poor SKCM patient prognosis (**Figure 1B**) and *SPINT1* expression was significantly inhibited in human SKCM comparing with nevus and normal skin (**Figure 1C**). We next performed a GO enrichment analysis of biological process (**Figure 1D**), analyzing the differentially expressed genes in SKCM samples of the TCGA cohort with missense mutations or deep deletion of *SPINT1*. The results showed that regulation of immune system, inflammatory response, cell cycle, cell adhesion, and extracellular matrix organization represent key pathways significantly affected in human SKCM with these *SPINT1* genetic alterations. Collectively, these results point to a role for *SPINT1* in SKCM aggressiveness and its crosstalk with the tumor immune microenvironment.

**Figure 1:**
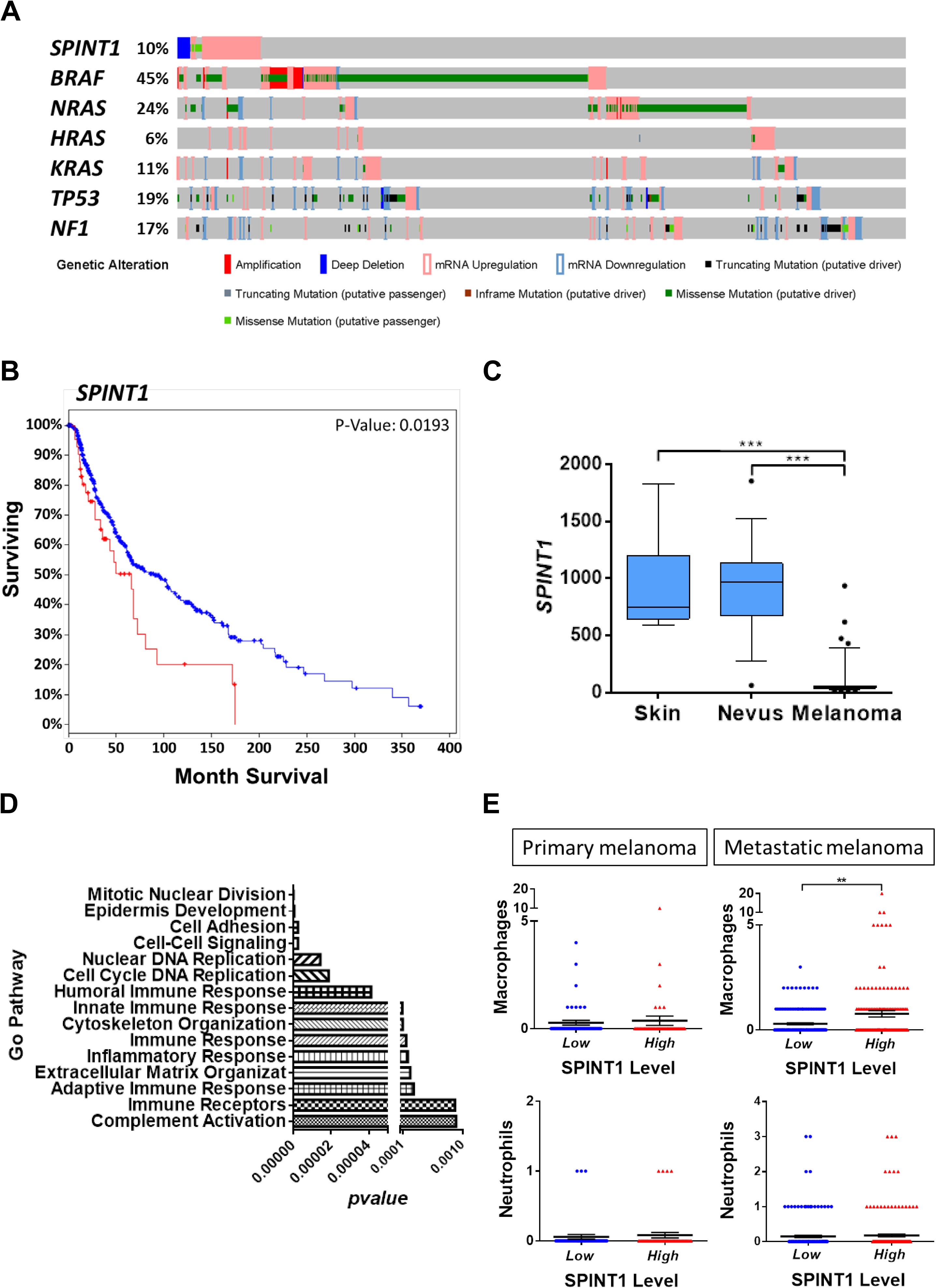
SPINT1 genetic alterations are associated with poor prognosis of SKCM patients. **(A)** Percentage of genetic alterations in oncogenes, tumor suppressor genes and *SPINT1* in SKCM patients of the TCGA cohort (n=479). **(B)** Survival curve of patients with genetic alteration (increased mRNA level, missense mutations and deep deletions) vs. wild type *SPINT1* of SKCM of the TCGA cohort. Kaplan–Meier Gehan-Breslow-Wilcoxon and nonparametric Log-rank Test. **(C)** Genetic expression of *SPINT1* in human samples from normal skin, nevus and malignant melanoma from GEO data set GDS1375 and 202826_at probe (n=70). ***p<0.001 according to ANOVA and Tukey’s Multiple Comparison Test. (D) Enrichment analysis of GO biological process. Representation of the most significant GO biological process altered when *SPINT1* is affected by missense mutations or deep deletion. Analysis Type: PANTHER Overrepresentation Test (Released 05/12/2017), Test Type: FISHER. **(E)** Number of infiltrated macrophages and neutrophils in SKCM samples of the TCGA cohort (n=479). The number of infiltrated cells in SKCM samples with low (blue) or high (red) *SPINT1* mRNA levels according to the median. The mean ± S.E.M. for each group is shown. *p<0.05; **p<0.01 according to Student *t* Tests.

The tumor microenvironment contains diverse leukocyte populations, including neutrophils, eosinophils, dendritic cells, macrophages, mast cells and lymphocytes (Coussens and Werb, 2002). It is known that tumor-associated macrophages (TAM) are able to interact with tumor cells and can promote cancer progression(Raposo et al., 2015). As shown in **Figure 1E**, the number of TAM in human SKCM samples correlated with the mRNA levels of *SPINT1* in metastatic SKCM. However, the number of tumor-associated neutrophils (TAN) was independent of *SPINT1* levels in both primary and metastatic SKCM. These data further confirmed the role of SPINT1 in the regulation of the crosstalk between tumor cells and inflammatory cells in human SKCM.

### The expression of SPINT1 positively correlates with both inflammation and macrophage markers in human SKCM biopsies

In order to further understand the role of SPINT1 in SKCM, the RNA Seq database of the large TCGA cohort of SKCM was analyzed in terms of the expression of *SOX10, TYR* and *DCT* genes, that have been shown to be important in melanocyte development (Ordonez, 2014; Ronnstrand and Phung, 2013). In addition, *SOX10* is a recognized biomarker for the diagnosis of SKCM (Ronnstrand and Phung, 2013). It was found that *SPINT1* expression positively correlated with those of *SOX10* and *TYR*, while a negative correlation was found between the expression of *SPINT1* and *DCT* (**Figure 2A**). The expression of the epithelial to mesenchymal transition (EMT) markers *ZEB1, ZEB2* and *TWIST1*, but not *TWIST2*, negatively correlated with that of *SPINT1* in SKCM (**Figure 2B**).

**Figure 2.**
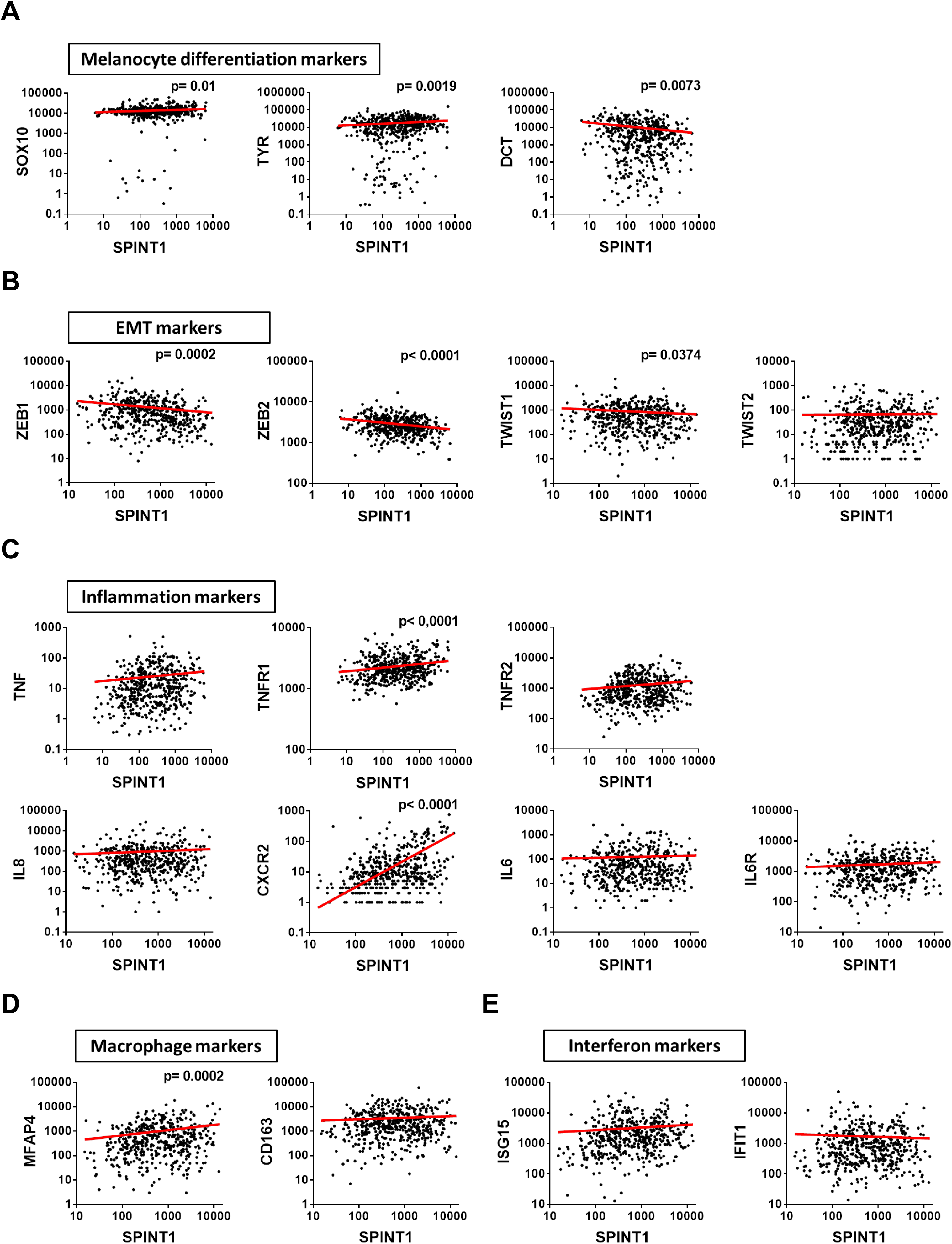
*SPINT1* expression correlates with aggressiveness marker expression in human SKCM biopsies. Correlation of *SPINT1* gene expression with those of the melanocyte differentiation markers *SOX10,TYR* and *DCT* **(A)** and the EMT markers *ZEB1,ZEB2,TWIST1* and *TWIST2* **(B)**, the inflammation markers *TNFA,TNFR1,TNFR2,IL8* (*CXCL8*), *CXCR2,IL6* and *IL6R* **(C)**, the macrophage markers *MFAP4* and *CD163* **(D)** and the interferon markers *ISG15* and *IFIT1* **(E)** in human SKCM biopsies of the TCGA cohort. The statistical significance of the correlation was determined using Pearson’s correlation coefficient. A linear regression-fitting curve in red is also shown.

SKCM cells release several cytokines and chemokines that recruit and polarize macrophages (Wang et al., 2017). Therefore, several inflammation markers were analyzed and only the expression of the genes encoding the receptor of the proinflammatory cytokine TNFα (TNFR1) and the receptor of the pro-inflammatory chemokine interleukin 8 (CXCR2), positively correlated with *SPINT1* levels (**Figure 2C**). Notably, the macrophage marker *MFAP4* also positively correlated with *SPINT1* expression (**Figure 2D**). However, the M2 polarization marker *CD163* (**Figure 2D**) and several interferon-stimulated genes (ISGs) (**Figure 2E**) were all unaffected by *SPINT1* levels. Collectively, these results further suggest that SPINT1 regulates SKCM differentiation and aggressiveness, and macrophages infiltration.

### Inflammation accelerates the onset of SKCM in zebrafish

Given the strong correlation between alterations of *SPINT1* levels with the progression of SKCM and the crosstalk with the tumor immune microenvironment, we crossed the zebrafish line *kita:Gal4;HRAS-G12V*, which expresses the human oncogene *HRAS-G12V* in melanocytes and spontaneously develops SKCM (Santoriello et al., 2010), with the zebrafish mutant line *spint1a*^*hi2217Tg/hi2217Tg*^ *(Mathias et al., 2007)*, which presents chronic skin inflammation (**Figure 3A**). Firstly, we quantified by fluorescence microscopy the number of early oncogenically transformed goblet cells, which also expressed the *kita* promoter(Feng et al., 2010), in *spint1a*-deficient larvae and their wild type siblings (**Figure 3B**). The results showed that *spint1a* deficiency resulted in increased number of HRAS-G12V^+^ cells (**Figure 3C**). To determine if the enhanced Spint1a deficiency-driven oncogenic transformation was also able to promote SKCM aggressiveness, SKCM development in *spint1a*^*hi2217Tg/hi2217Tg*^ fish were compared with wild type (*spint1a*^*+/hi2217Tg*^) from the end of metamorphosis stage (between 28-30 dpf) to 120 dpf (adult stage) (**Figures 3D-3F**). The resulting Kaplan-Meier curve showed a significant decreased tumor-free rate in the Spint1a-deficient fish, which developed SKCM in more than 50% of cases at 50-60 dpf compared with their wild type siblings which reached only 75% at this age (**Figure 3F**). These data suggest that Spint1a deficiency increases oncogenic transformation and accelerates SKCM onset *in vivo*.

**Figure 3:**
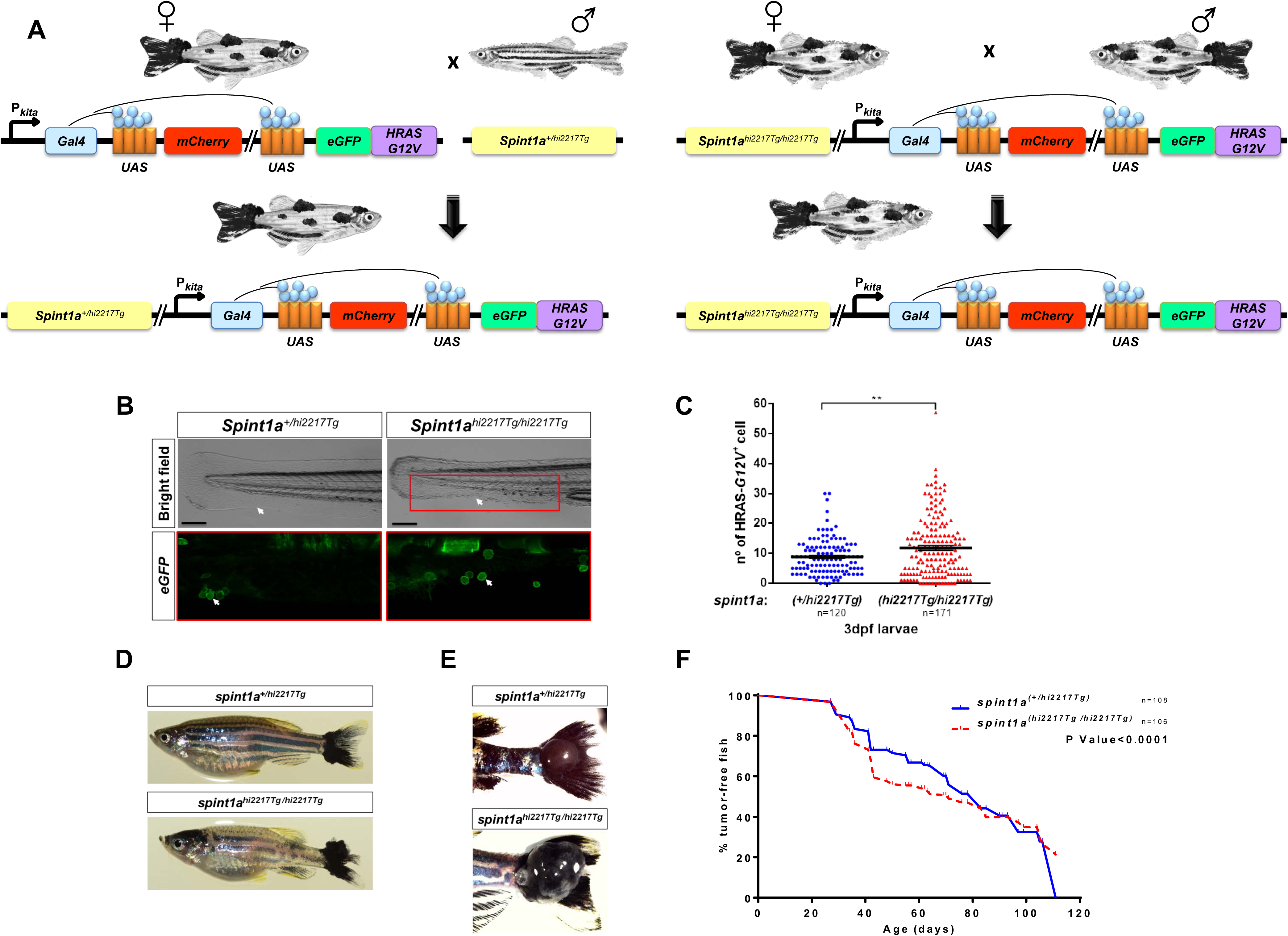
Inflammation accelerates the onset of SKCM in zebrafish. **(A)** Schematic diagram of the generation of the SKCM model line in zebrafish with Spint1a deficiency. **(B-C)** Representative images (B) and number of early oncogenically transformed eGFP-HRAS-G12V^+^ cells in the boxed area (C) in Spint1a-deficient larvae and control siblings at 3 dpf. Note the morphological alterations observed in the inflamed skin of the mutants (white arrows). eGFP-HRAS-G12V^+^ goblet cells are marked with white arrows. Scale bar 250 µm. Each point on the scatter plot represents one larva and the mean ± SEM is also shown. ** p<0.05 according to an unpaired Student *t* test with Welch’s correction. **(D-F)** Impact of Spint1a deficiency on SKCM onset in zebrafish. Representative images of whole fish **(D)** and of nodular tail tumors **(E)**, and Kaplan-Meier curve showing the percentage of SKCM-free fish in control and Spint1a-deficient adult fish **(F)**. p < 0.0001 according to a Log rank Mantel-Cox test.

### Spint1a deficiency is required at cell autonomous and non-autonomous levels to enhance SKCM cell dissemination in a zebrafish larval allotrasplantation model

To assess the *in vivo* role of Spint1a deficiency in SKCM invasiveness, SKCM tumors from *spint1a*^*hi2217Tg/hi2217Tg*^; *kita:Gal4;HRAS-G12V* and *spint1a*^*+/hi2217Tg*^; *kita:Gal4;HRAS-G12V* were disaggregated, after staining the cells were transplanted into the yolk sac of 2 dpf casper zebrafish larvae (**Figure 4A**). The results showed that Spint1a deficiency in SKCM cells enhanced the dissemination of SKCM, assayed as the percentage of invaded larvae and the number of foci per larva, compared to control SKCM cells (**Figures 4B-4D**). We next examined whether Spint1a deficiency in the stroma, i.e. in a non-autonomous manner, also promoted SKCM aggressiveness. Spint1a wild type SKCMs were transplanted into the yolk sac of Spint1a-deficient and their wild type siblings larvae (**Figure 4E**). Strikingly, it was found that Spint1a deficiency in the tumor microenvironment also promoted a significantly higher dissemination of SKCM compared to control tumor microenvironments (**Figures 4F-4H**).

**Figure 4:**
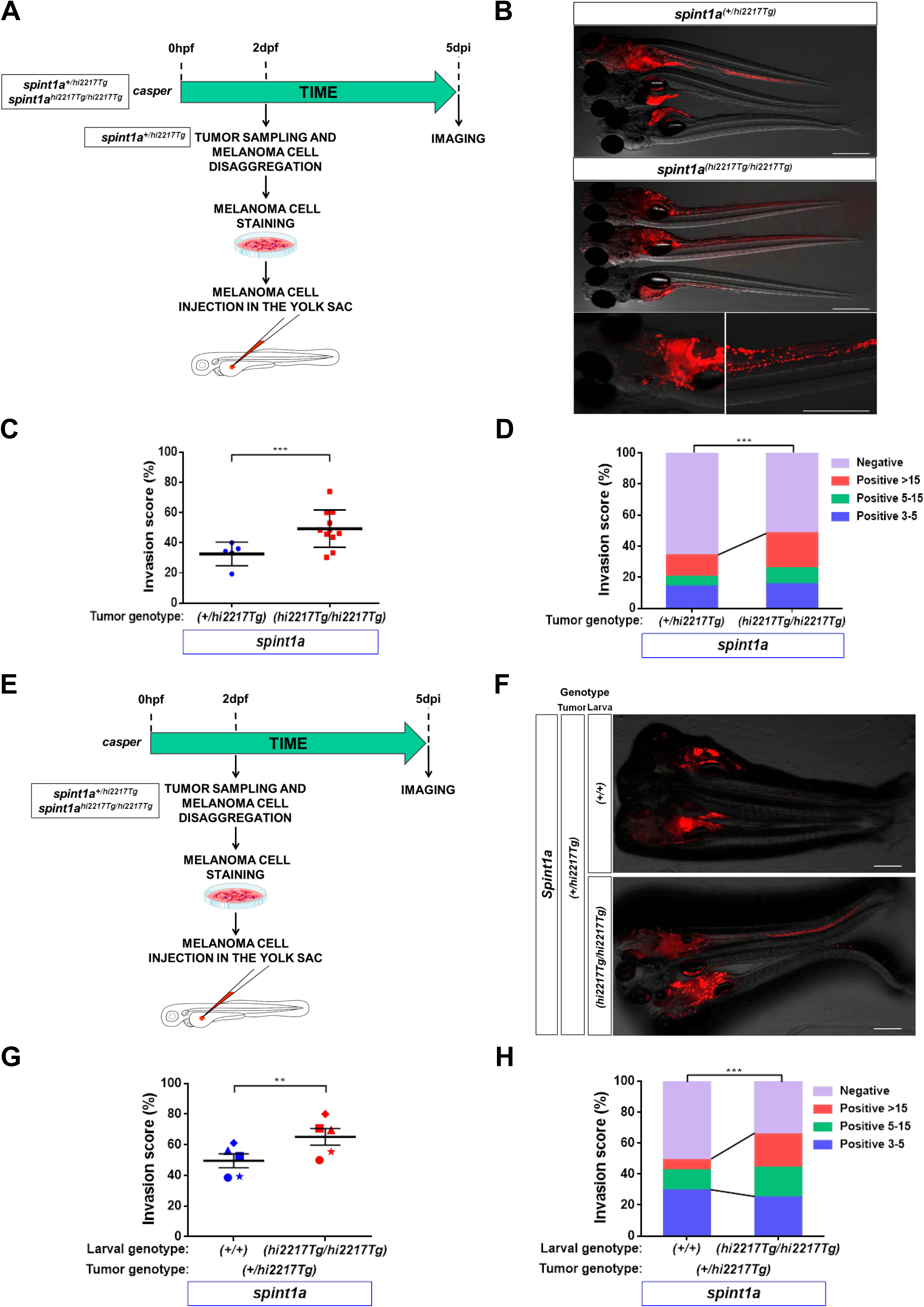
Spint1a deficiency is required at cell autonomous and non-autonomous levels to enhance SKCM cell dissemination in a zebrafish larval allotrasplantation model. Analysis of dissemination of control and Spint1a-deficient SKCM allotransplants in wild type larvae **(A-D)** and SKCM allotransplants in wild type and Spint1a-deficient larvae **(E-H)**. **(A,E)** Experimental design. **(B,F)** Representative images (overlay of bright field and red channels) of SKCM invasion at 5 dpi. Bars: 500 µm. **(C,G)** Percentage of invaded larvae for both tumor genotypes. Each dot represents a single tumor and the mean ± SEM is also shown. **p<0.01, ***p< 0.0001 according to unpaired Student *t* test. (D, H) Number of tumor foci per larva. ***p< 0.0001 according to Chi-square Tests.

To further confirm a role of Spint1a in both SKCM and tumor microenvironment cells, we next sorted tumor (eGFP^+^) and stromal (eGFP^−^) cells from both genotypes and then mixed in equal proportions (∼90% of tumor and ∼10% of stromal cells) in the 4 possible combinations (**Figure 5A**), since it was found that all tumors had ∼90% of tumor and ∼10% of stromal cells (data not shown). Notably, both Spint1a-deficient tumor and stromal cells were able to increase SKCM cell invasion (**Figure 5B and 5C**). Collectively, these results suggest that Spint1a deficiency enhances SKCM invasion by both cell autonomous and non-autonomous mechanisms.

**Figure 5:**
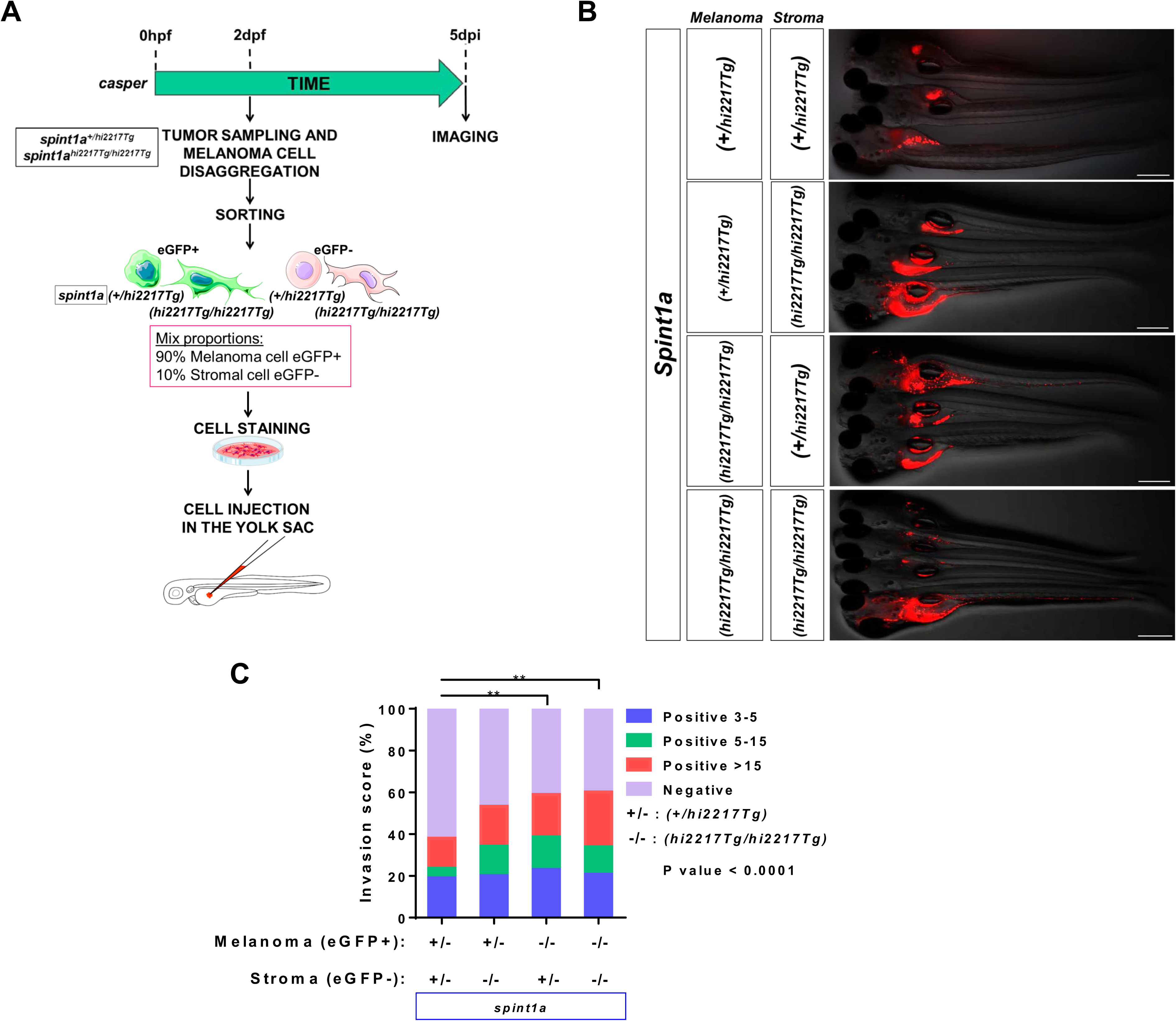
Spint1a deficiency both in stromal and tumor cell enhances SKCM dissemination in zebrafish larval model. **(A)** Allotransplant experimental design. Combinations of Spint1a-deficient tumor and stromal cells from SKCMs were mixed with wild type tumor and stromal cells. All possible cell combinations were obtained maintaining the initial ratio. **(B)** Representative images (overlay of bright field and red channels) of the invasion in wild type recipient larvae at 5 dpi. Bar: 500 µm. **(C)** Number of tumor foci per larva. **p< 0.01 according to a Chi-square Test.

### Spint1a-deficient SKCM cells showed enhanced aggressiveness in adult zebrafish allotransplantation model

The results obtained in allotransplantation assay in larvae prompted us to analyze the role of Spint1a in SKCM aggressiveness and metastasis in adult casper zebrafish to directly visualize tumor cell proliferation and dissemination over time. *spint1a*^*hi2217Tg/hi2217Tg*^ and *spint1a*^*+/hi2217Tg*^ SKCMs were sampled, disaggregated and subcutaneously injected (300,000 cells) in the dorsal sinus of adult casper recipients previously irradiated with 30 Gy (**Figure 6A**). Tumor engraftment was visible as early as 7 days post-transplantation in both genotypes. While 90% engraftment was obtained with wild type SKCM cells, Spint1a-deficient cells showed a significant enhancement of tumor engraftment rate, around 95% (**Figure 6B**). In addition, adult zebrafish recipients transplanted with Spint1a-deficient SKCMs developed tumors with a significant higher growth rate than those injected with wild type SKCMs (**Figures 6C**). Notably, Spint1a-deficient SKCM cells were able to invade the entire dorsal area, part of ventral cavity and the dorsal fin (**Figures 6C**).

**Figure 6:**
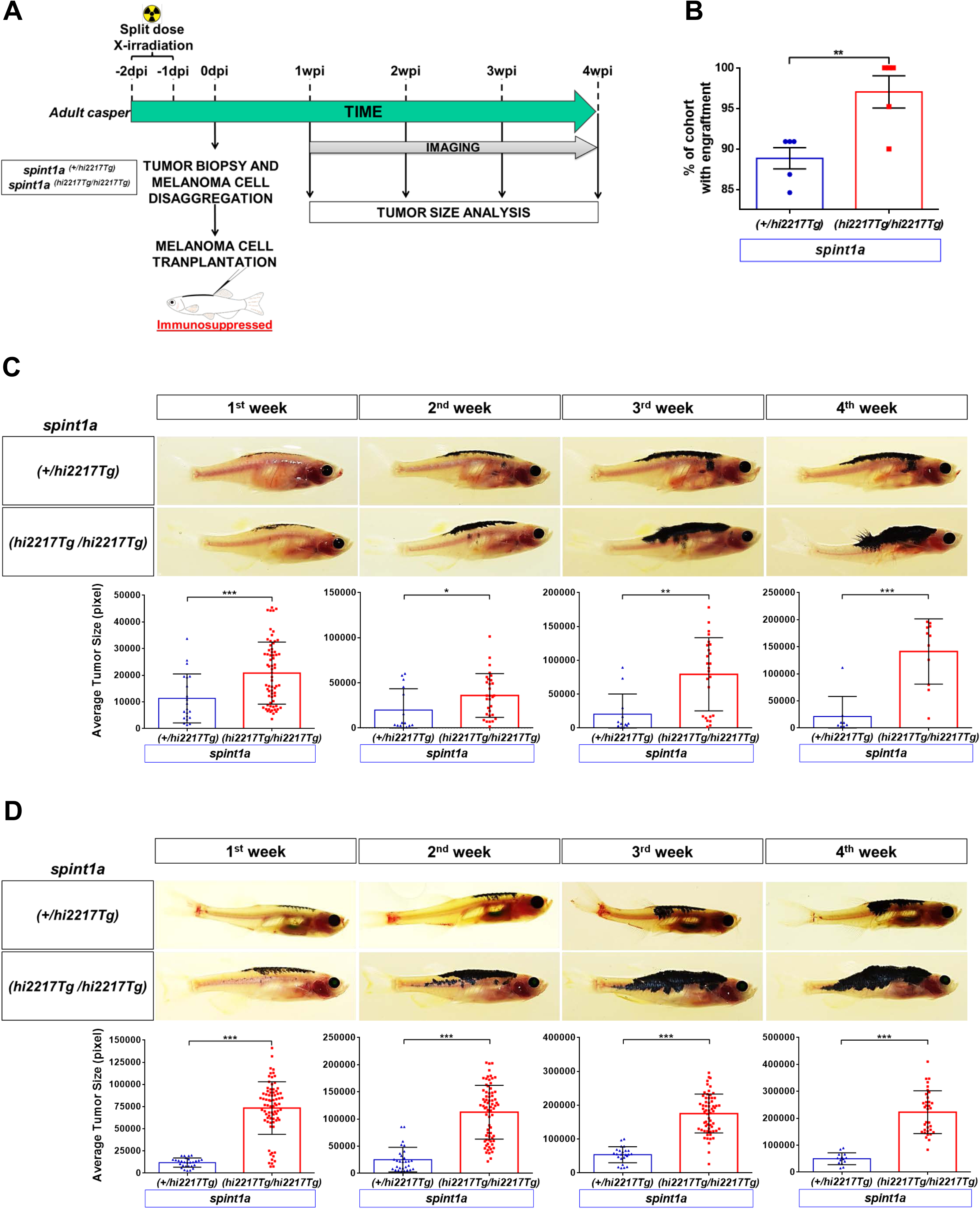
Spint1a-deficient SKCM shows enhanced aggressiveness in adult zebrafish allotransplantation assays. **(A)** Experimental workflow of adult allotransplantation experiments in pre-irradiated adult casper zebrafish. **(B)** Percentage of engraftment for both control and Spint1a-deficient tumors. Each dot represents a single SKCM tumor and the mean ± SEM is also shown. **(C,D)** Representative images and average tumor size (pixels) from 1 to 4 weeks post-transplant of primary **(C)** and secondary **(D)** transplants. Each dot corresponds to a recipient-transplanted fish and the mean ± SEM is also shown. **(B-D)** *p < 0.05, **p<0.01, ***p<0.001 according to unpaired Student *t* test.

We next performed additional transplant assays following the same work-flow but injecting an increased number of cells (500,000 cells per recipient fish), that ensured a 100 % of engraftment was for both genotypes (data not shown). From the first week of analysis, Spint1a-deficient SKCM tumor size was significantly larger than their control counterparts (**Figure 6D**). In addition, the recipients injected with Spint1a-deficient SKCM cells developed larger tumors spanning the entire dorsal area and even exceed the notochord line and grew vertically, a clear aggressiveness signature of SKCM (**Figure 6D**).

To further investigate the aggressiveness potential of Spint1a-deficient SKCMs, a serial dilution assay was performed following the work-flow previously described in **Figure S1**. Cells from both Spint1a-deficient and wild type SKCMs were serially diluted and 3 different numbers of cells (30,000 cells, 100,000 cells and 300,000 cells) were transplanted into recipients as described above. Notably, while 30,000 and 100,000 Spint1a-deficient SKCM cells were able to engraft and the tumor grew over the time, wild type SKCMs hardly grew (**Figures S1A, S1B**). However, injection of 300,000 Spint1a-deficient SKCM cells resulted in large tumor spanning the entire dorsal area and invading part of the ventral cavity (**Figure S1C**), confirming previous results. Collectively, all these results confirm that Spint1a deficiency enhances SKCM aggressiveness.

### Spint1a deficiency promotes SKCM dedifferentiation and inflammation

To understand the mechanisms involved in the Spint1a-mediated aggressiveness of SKCM, the expression of genes encoding important biomarkers was analyzed by RT-qPCR. The mRNA levels of *sox10, tyr, dct* and *mitfa* were lower in Spint1a-deficient SKCMs than in their wild type counterparts (**Figure S2**). In addition, while the transcript levels of *mmp9 and slug* were similar in Spint1a-deficient and wild type SKCM, *cdh1* levels were significantly decreased in Spint1a-deficient compared to wild type SKCM (**Figure S2**).

We next analyzed genes encoding key inflammatory molecules and immune cell markers, including the pro-inflammatory cytokine Il1b, the neutrophil markers Lyz and Mpx, the macrophage marker Mpeg1, and the ISGs B2m, Mxb and Pkz, in Spint1a-deficient and wild type SKCMs (**Figure S2**). Curiously, it was found that while *il1b, lyz* and *mpx* mRNA levels were not affected by Spint1a deficiency, those of *mpeg1* were elevated in Spint1a-deficient SKCMs. Furthermore, the ISGs *b2m, mxb* and *pkz* genes showed enhanced mRNA levels in Spint1a-deficient SKCMs. These results point out to altered immune surveillance and tumor cell dedifferentiation promoted by Spint1a-deficiency in SKCM.

## Discussion

The relationships between inflammation and cancer are ambiguous. Although it is estimated that underlying infections and inflammatory responses are linked to 15– 20% of all deaths from cancer worldwide (Maru et al., 2014), immunosuppression is known to increase the risk for certain tumors (Shalapour and Karin, 2015). Furthermore, immunotherapy is considered the most promising cancer therapy for the next future (Carreau and Pavlick, 2019). In this study, we have developed a preclinical zebrafish model to study the role of SPINT1-driven skin chronic inflammation in melanoma. We found that Spint1a deficiency is required at both cell autonomous and non-autonomous levels to enhance cell dissemination of SKCM by promoting tumor dedifferentiation and altered immune surveillance. These results may have important clinical impact, since genetic alterations of SPINT1 were found in 10% of SKCM patients and correlated with altered cell cycle, differentiation and innate and adaptive immune signaling pathways and, more importantly, with a poor prognosis. In addition, *SPINT1* transcript levels positively correlated with macrophage infiltration, but not neutrophil one, in SKCM tumor samples. Curiously, activated neutrophils in a condition of repeated wounding have been shown to interact with pre-neoplastic cells promoting their proliferation through the release of prostaglandin E2 and, more importantly, SKCM ulceration correlates with increased neutrophil infiltration and tumor cell proliferation, which are both associated with poor prognosis (Antonio et al., 2015). Although we found a robust positive correlation between the transcript levels of *SPINT1* and *CXCR2*, which encodes a major IL-8 receptor involved in SKCM neutrophil infiltration (Jablonska et al., 2014), CXCR2 has also been shown to promote tumor-induced angiogenesis and increased proliferation(Gabellini et al., 2018; Gabellini et al., 2009; Singh et al., 2009) and, therefore, the SPINT1/CXCR2 axis may regulate SKCM aggressiveness by neutrophil-independent pathways.

The zebrafish model developed in this study uncovered a role for Spint1a in facilitating oncogenic transformation which probably accelerates the SKCM onset. Although SPINT1 is a serine protease inhibitor with several targets, including ST14 and HGFA, deregulation of the SPINT1/ST14 axis leads to spontaneous squamous cell carcinoma in mice (List et al., 2005) and keratinocyte hyperproliferation in zebrafish (Carney et al., 2007; Mathias et al., 2007) preceded by skin inflammation in both models. In addition, intestine-specific Spint1 deletion in mice induces the activation of the master inflammation transcription factor NF-κB and accelerated intestinal tumor formation (Kawaguchi et al., 2016). Strikingly, pharmacological inhibition of NF-κB activation reduces the formation of intestinal tumors in Spint1-deficient ApcMin/+ mice(Kawaguchi et al., 2016), unequivocally demonstrating that Spint1-driven inflammation promotes tumorigenesis.

The SKCM allotransplant assays in larvae revealed for the first time that Spint1a deficiency in both tumor and stromal cells increases SKCM invasiveness. In addition, Spint1a deficiency in both cell types does not show enhanced invasiveness compared to Spint1a deficiency in either cell type. This is an interesting observation, since SPINT1 is a membrane-bound protein that may, therefore, inhibit their targets in both tumor cell autonomous and non-autonomous manners. However, wild type Spint1a tumor microenvironment fails to compensate its loss in tumor cells, since Spint1a-deficient tumor cells show enhanced invasiveness in wild type larvae and adult recipients, and vice versa. Importantly, transplantation experiments of serial diluted SKCM cells revealed the crucial cell-autonomous role of Spint1a in inhibiting tumor aggressiveness. Similarly, loss of SPINT1 in human pancreatic cancer cells promotes ST14-dependent metastasis in nude mouse orthotopic xenograft models (Ye et al., 2014). We observed that genetic alterations in *SPINT1* transcript levels in SKCM patient samples negatively correlated with EMT markers and that Spint1a-deficient zebrafish SKCM showed reduced *cdh1* mRNA levels. EMT phenotype switching has been shown to be involved in acquisition of metastatic properties in the vertical growth phase of SKCM (Bennett, 2008) and loss of E-cadherin, with gain of N-cadherin and osteonectin, was associated with SKCM metastasis(Alonso et al., 2007). Importantly, the presence of aberrant E-cadherin expression in primary and metastatic SKCM is associated with a poor overall patient survival(Yan et al., 2016). Therefore, our results suggest that SPINT1 loss may facilitate metastatic invasion of human SKCM through EMT phenotype switching.

In summary, we have developed a preclinical model to study the role of altered expression of SPINT1 in early transformation, progression and metastatic invasion in SKCM. This model has revealed that Spint1a deficiency facilitates oncogenic transformation, regulates the tumor/immune microenvironment crosstalk and is associated to SKCM aggressiveness. In addition, Spint1a deficiency in either tumor or microenvironment compartment increases SKCM aggressiveness. The high prevalence of *SPINT1* genetic alterations in SKCM patients and their association with a poor prognosis, suggest the relevance of clinical intervention on this signaling pathway for precision SKCM medicine.

## Materials and Methods

### Animals

The experiments complied with the Guidelines of the European Union Council (Directive 2010/63/EU) and the Spanish RD 53/2013. Experiments and procedures were performed as approved by the Consejería de Agua, Agricultura, Ganadería y Pesca de la CARM (authorization number # A13180602).

Wild-type zebrafish (*Danio rerio* H. Cypriniformes, Cyprinidae) were obtained from the Zebrafish International Resource Center (ZIRC, Oregon, USA) and mated, staged, raised and processed as described in the zebrafish handbook (Westerfield, 2000). Zebrafish fertilized eggs were obtained from natural spawning of wild type and transgenic fish held at our facilities following standard husbandry practices. Animals were maintained in a 12 h light/dark cycle at 28°C. *Tg(kita:GalTA4,UAS:mCherry)*^*hzm1*^ zebrafish were crossed with *Tg(UAS:eGFP-H-RAS_G12V)*^*io6*^ line (Santoriello et al., 2010) to express oncogenic human HRAS_G12V driven by the melanocyte cell-specific promoter *kita*. The hi2217 line, which carries a hypomorphic *spint1a* mutant allele that promotes skin inflammation (Mathias et al., 2007), and transparent *roy*^*a9/a9*^; *nacre*^*w2/w2*^ (casper)(White et al., 2008) of 4-8 month old were previously described.

Zebrafish larvae were anesthetized by a solution of 0.16 mg/ml buffered tricaine (Sigma-Aldrich) in embryo medium. Adult zebrafish were anesthetized by a dual anesthetic protocol to minimize over-exposure to tricaine, in long-term studies (up to 40 min) (Dang et al., 2016). Briefly, the anesthesia was firstly induced by tricaine and then fish were transferred to tricaine/isoflurane solution (forane in ethanol, 1:9).

### Tumor sampling, disaggregation and cell sorting

Primary melanoma tumors were excised from adult zebrafish once they had reached between 3-5 mm in diameter. Some individuals were euthanized according the European Union Council and IUAC protocol and others were monitored and maintained still alive after the tumor biopsy treated with conditioners to reduce fish stress and heal damaged tissue and wounds (STRESS COAT, API), as well as to protect from bacterial (MELAFIX, API) and fungal infections (PIMAFIX, API).

The tumor was excised with a clean scalpel and razor blade, placed in 2 ml of dissection media, composed by DMEM/F12 (Life Technologies), 100 UI/ml penicillin, 100 µg/ml streptomycin, 0.075 mg/ml Liberase (Roche). After manually disaggregation with a clean razor blade and incubation at room temperature for 30 min, 5 ml of wash media, composed by DMEM/F12 (Life Technologies), penicillin-streptomycin (Life Technologies), and 15% heat-inactivated FBS (Life Technologies), were added to the tumor slurry and manually disaggregated one last time. Next, the tumor cell suspensions were passed through a 40 µm filter (BD) into a clean 50 ml tube. An additional 5 ml of wash media was added to the initial tumor slurry and additionally filtered. This procedure was repeated twice. Cells were counted with a hemocytometer and the tubes of resuspended cells were centrifuged at 500 g for 5 min. The pellets of tumor cells were resuspended in PBS containing 5% FBS and kept on ice prior to transplantation (Dang et al., 2016).

The resulting cell suspension from zebrafish melanoma tumors was passed through a 40 μm cell strainer and propidium iodide (PI) was used as a vital dye to exclude dead cells. The Cell Sorting was performed on a “Cell Sorter” SONY SH800Z in which eGFP positive cells were sorted from the negative ones of the same cell tumor suspension.

### Larval allotransplantion assays

Melanomas were disaggregated, then labelled with 1,1’-di-octa-decyl-3,3,3’,3’-tetra-methyl-indo-carbo-cya-nine perchlorate (DiI, ThermoFisher) and finally resuspended in a buffer containing 5% FBS in PBS. Between 25 to 50 cells/embryo were then injected in the yolk sac of Casper or *spint1* mutants zebrafish larvae 48 hours post-fertilization (hpf) and after 5 days at 28°C, larvae were analyzed for zebrafish melanoma cells dissemination by fluorescence microscopy (Marques et al., 2009). Melanoma cell invasion score was calculated as the percentage of zebrafish melanoma cell-invaded larvae over the total number of larvae analyzed taking into account also the number of tumor foci per larvae. Three tumor foci were established to score a larva as positive for invasion. Furthermore, larvae positive for invasion were also distinguished in three groups considering the number of positive foci per larvae: 3-5 foci per larvae, 5-15 foci per larvae and >15 foci per larvae.

### Adult allotransplantion assays

Adult zebrafish used as transplant recipients were immunosuppressed to prevent rejection of the donor material. Thus, the recipients were anesthetized, using the dual anesthetic protocol described above, and treated with 30 Gy of split dose sub-lethal X-irradiation (YXLON SMART 200E, 200 kV, 4.5 mA) two days before the transplantation. Then the immunosuppressed fish were maintained in fresh fish water treated with conditioners preventing infections onset and the consequent recipient deaths.

Anesthetized fish were placed dorsal side up on a damp sponge and stabilized with one hand. Using the other hand, the needle of a 10 µl beveled, 26S-guaged syringe (Hamilton) was positioned midline and ahead to the dorsal fin. 30,000, 100,000, 300,000 and 500,000 cells resuspended in PBS were injected into the dorsal subcutaneous cavity. The syringe was washed in 70% ethanol and rinsed with PBS between uses. Following transplantation, fish were placed into a recovery tank of fresh fish water and kept off-flow with daily water changes for 7 days. Large and pigmented tumors engrafted and were observed to expand by 10 days post-transplantation.

### SKCM imaging in adult zebrafish

Adult zebrafish were scored weekly for melanoma formation starting at the first appearance of raised lesions. Tumor scoring was blinded and experiments were independently repeated at least 3 times. Zebrafish were anesthetized, placed in a dish of fish water and photographed using a mounted camera (Nikon D3100 with a Nikon AF-S Micro Lens). The pigmented tumor size was represented as the number of pigmented pixels (Adobe Photoshop CS5).

### Analysis of gene expression

Once zebrafish tumors reached between 3-5 mm of diameter, they were excised and total RNA was extracted with TRIzol reagent (Invitrogen), following the manufacturer’s instructions, and then treated with DNase I, amplification grade (1 U/µg RNA; Invitrogen). SuperScript III RNase H^−^ Reverse Transcriptase (Invitrogen) was used to synthesize first-strand cDNA with oligo(dT)_18_ primer from 1 µg of total RNA at 50°C for 50 min. Real-time PCR was performed with an ABI PRISM 7500 instrument (Applied Biosystems) using SYBR Green PCR Core Reagents (Applied Biosystems). Reaction mixtures were incubated for 10 min at 95°C, followed by 40 cycles of 15 s at 95°C, 1 min at 60°C, and finally 15 s at 95°C, 1 min 60°C and 15 s at 95°C. For each mRNA, gene expression was normalized to the ribosomal protein S11 (rps11) content in each sample Pfaffl method (Pfaffl, 2001). The primers used are shown in Table S1. In all cases, each PCR was performed with triplicate samples and repeated at least in two independent samples.

### Human SKCM dataset analysis

Normalized gene expression, patient survival data, genetic alterations and neutrophil/macrophage infiltration were downloaded from SKCM repository of The Cancer Genome Atlas (TCGA, Provisional) from cBioPortal database (https://www.cbioportal.org/). Transcript levels of *SPINT1* in human samples from normal skin, benign and malignant melanoma was collected from Gene Expression Omnibus (GDS1375 dataset and 202826_at probe). Gene expression plots and regression curves for correlation studies were obtained using GraphPad Prism 5.03 (GraphPad Software).

### Statistical analysis

Data are shown as mean ± SEM and they were analyzed by analysis of variance (ANOVA) and a Tukey multiple range test to determine differences between groups. The survival curves were analysed using the log-rank (Mantel-Cox) test. All the experiments were performed at least three times, unless otherwise indicated. The sample size for each treatment is indicated in the graph and/or in the figure legend. Statistical significance was defined as p < 0.05.

## Acknowledgments

We strongly acknowledge P. Martínez for his excellent technical assistance with zebrafish maintenance.

## Author contributions

VM conceived the study; EGA, LIZ, CG and VM designed research; EGA, SIM, DGM, IF and CG performed research; EGA, SIM, DGM, IF, LIZ, MCM, MLC, CG and VM analyzed data; and EGA, CG and VM wrote the manuscript with minor contribution from other authors.

---

**Figure S1.**
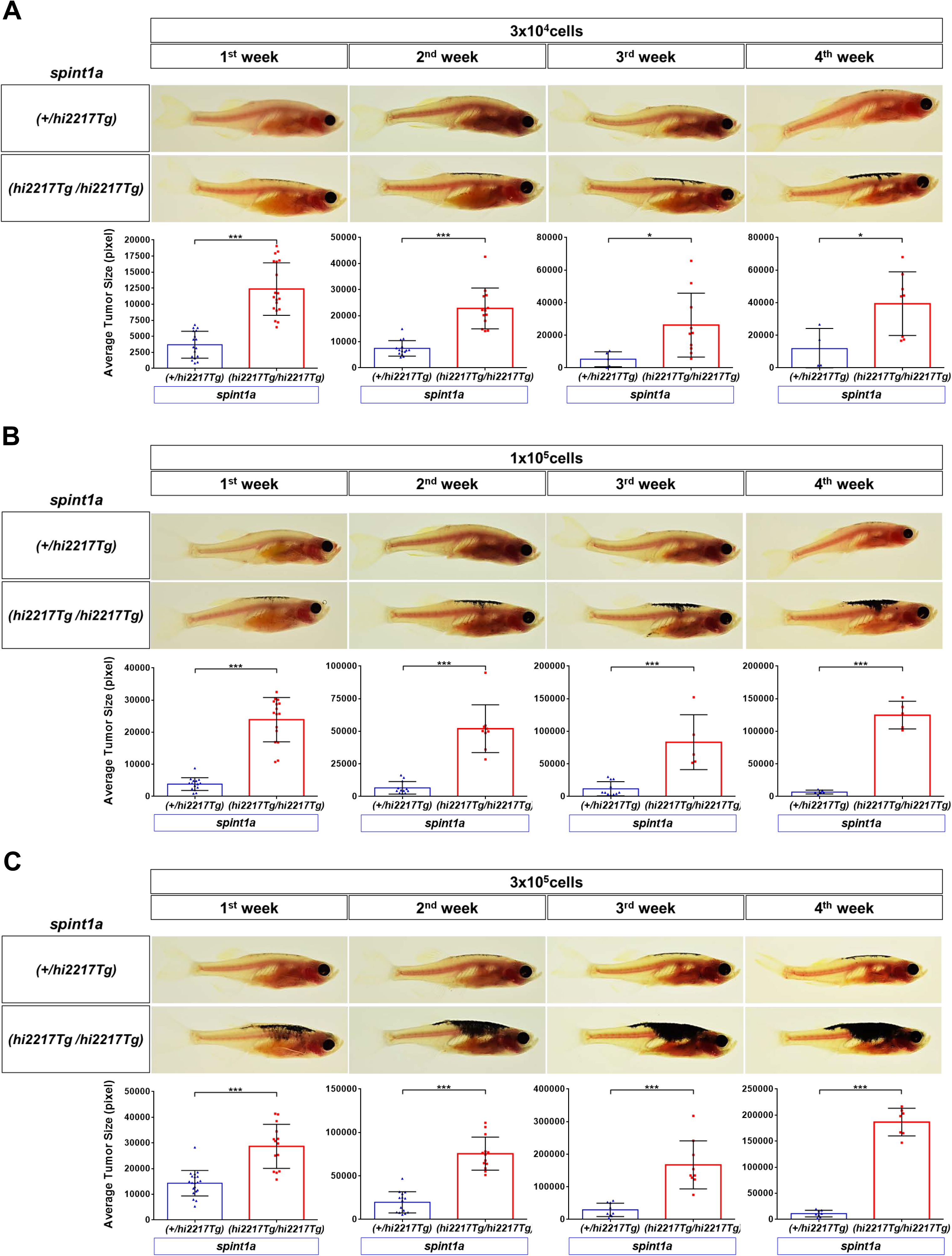
Spint1a-deficient SKCM shows enhanced aggressiveness in adult zebrafish allotransplantation assays. Control and Spint1a deficient SKCMs were disaggregated and 30,000 **(A)**, 100,000 **(B)** and 300,000 cells **(C)** were injected subcutaneously in pre-irradiated adult casper zebrafish. Fish were analyzed for average tumor size (pixels) from 1 to 4 weeks post-transplant. Representative images and quantification of the average tumor size are shown. Each dot corresponds to a recipient-transplanted fish and the mean ± SEM is also shown.*p<0.05, ***p<0.001 according to unpaired Student *t* test.

**Figure S2.**
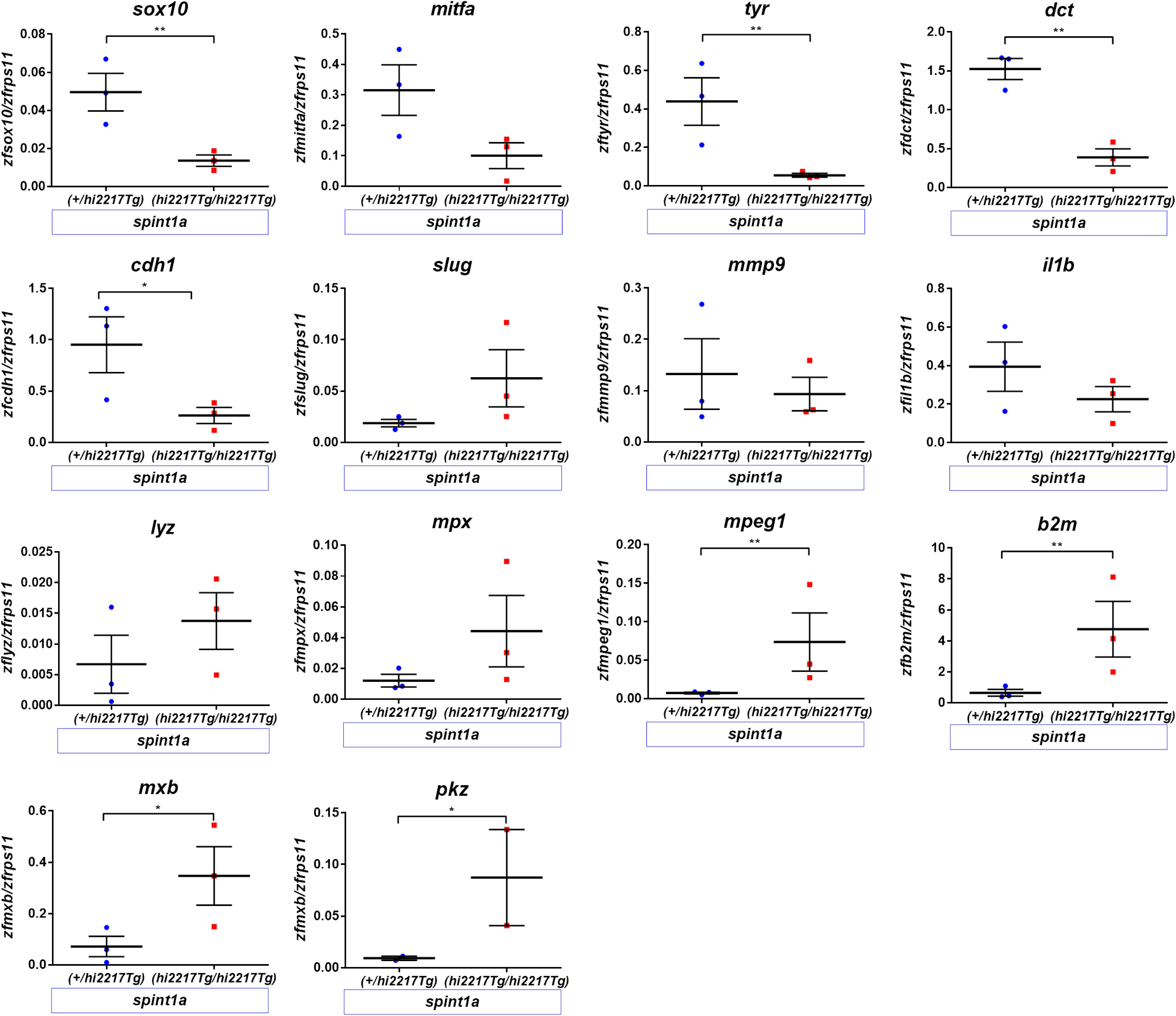
Expression analysis of differentiation melanocyte, EMT, inflammation and immune cell markers in zebrafish SKCM. The mRNA levels of the genes encoding the differentiation melanocyte markers *Sox10, Mitfa, Tyr* and *Dct*, the EMT markers *Cdh1, Slug* and *Mmp9*, the inflammation marker *Il1b*, the neutrophil markers *Lyz* and *Mpx*, the macrophage marker *Mpeg1*, and the ISGs *B2m, Mxb* and *Pkz* were analyzed by RT-qPCR in control and Spint1a-deficient SKCMs. *p < 0.05, **p<0.01 according to one-tailed Student *t* test.

**Table S1.**
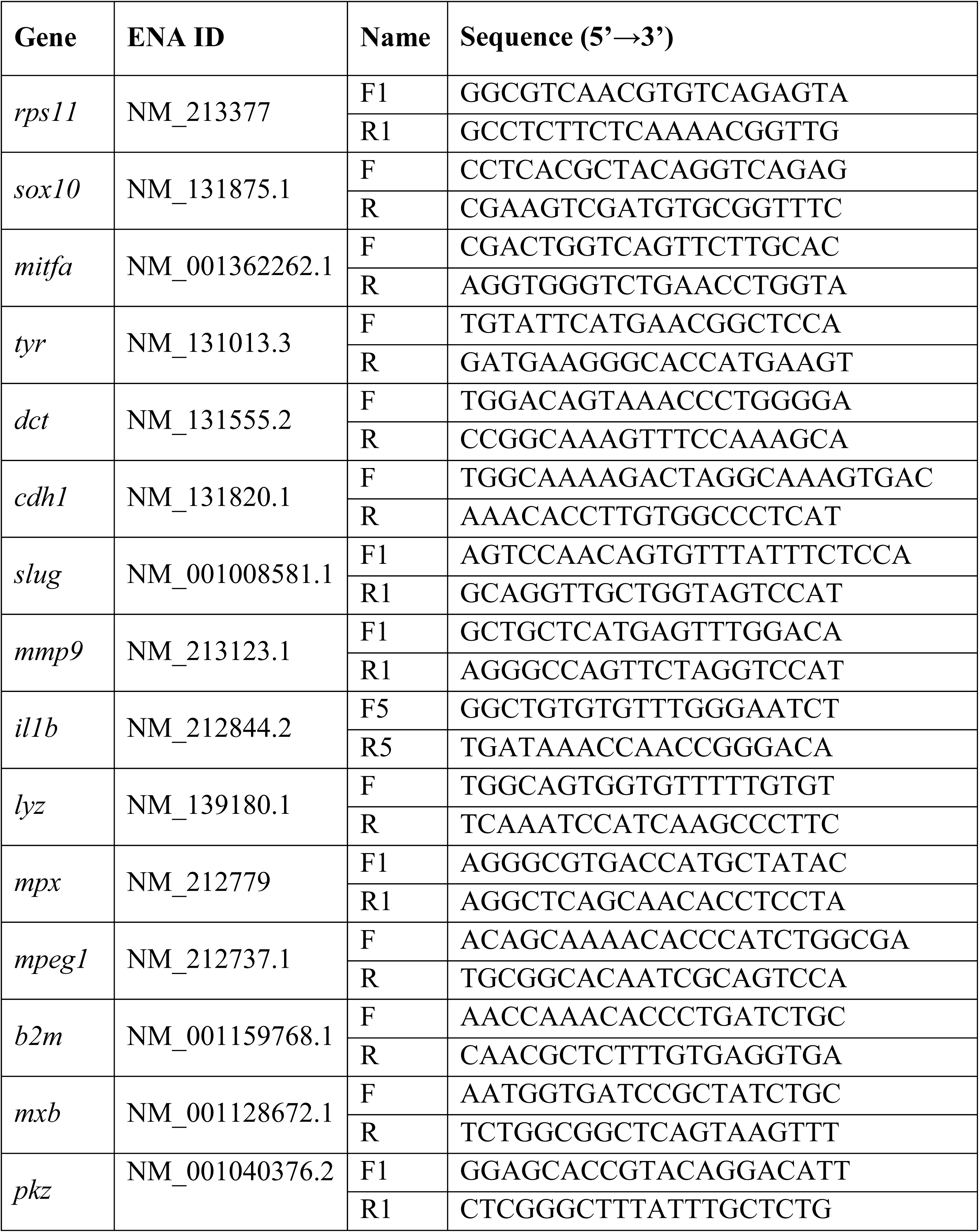
Primers used in this study for RT-qPCR. The gene symbols followed the Zebrafish Nomenclature Guidelines (http://zfin.org/zf_info/nomen.html/). ENA, European Nucleotide Archive (http://www.ebi.ac.uk/ena/).

